# Entomopoxvirus-like long DNA sequences in human centromeric and peri-centromeric regions

**DOI:** 10.1101/2024.11.07.622552

**Authors:** Eiichi Hondo, Tetsuya Mizutani, Hiroshi Shimoda, Atsuo Iida

## Abstract

Mammalian genomes harbor a range of endogenous virus-like sequences, although these sequences are typically affected by mutations and deletions, and no examples have been reported of long, conserved regions extending tens of kilobases or more. In this study, a comprehensive similarity search against the complete human T2T genome (hT2T) using high-quality viral genome data was performed. Among nearly continuous viral sequences, the longest identified was *Entomopoxvirus*, spanning 140 kb with a similarity of over 57%. Notably, *Entomopoxvirus*-like sequences stood out within the human genome, with a total of 2.42 Mb exhibiting similarity to human sequences. Seven *Entomopoxvirus* species were fragmented into regions less than 40 kb each, with nearly all of these segments scattered throughout the human genome. Surprisingly, nearly all of these sequences were localized within the hsat1A regions of human centromeric and pericentromeric areas. On chromosome 3, for example, all sequences were confined within the centromere, and extensive transcription was observed across these regions. While the lack of fully sequenced genomes with solid annotations for other eukaryotic species limits our ability to infer the phylogenetic significance of these regions, our findings reveal remarkably long virus-like sequences within centromeric and related regions—as integral components of eukaryotic chromosomes.

## Introduction

Our genomes contain various virus-like sequences known as endogenous viral elements (EVEs) [1, 2]. These sequences often exhibit mutations and deletions compared to intact viral genomes, making it rare to find long, complete viral sequences. However, it has long been recognized that EVEs play a critical role in regulating gene expression and can function as enhancers, making them significant for biological functions [3, 4]. Open chromatin regions, in particular, are considered more susceptible to the integration of foreign viral genomes, a mechanism which might enable retroviral EVEs to act as enhancers near host genes. Indeed, chimeric sequences between host genomes and HIV genomes have been observed in B cells of HIV-infected individuals, suggesting that these regions might serve as enhancer elements [5]. For these sequences to be passed to offspring, however, they must integrate through germline cells.

In mammals, retroviral EVEs have been co-opted as functional genes crucial for specific adaptations, such as PEG10 and syncytin, which are essential for placental development [6, 7], and SASPase, which is implicated in skin hydration [8]. A homolog closely related to the VP35 protein of Ebola and Marburg viruses has also been identified in *Myotis* bats, potentially linking these non-retroviral EVEs to the species’ ability to tolerate infections by these viruses without disease symptoms [9]. Retroviral EVEs, also known as endogenous retroviruses, constitute about 9% of our genome and are recognized as important viral-like sequences from an evolutionary perspective. For instance, endogenized viruses have been shown to influence coat color phenotypes in mice [10, 11]. Furthermore, retroviral EVEs in pigs have raised public health concerns due to their ability to release viral particles and to infect human cells [12]. Also, recent reports reveal that in certain bat species, approximately 30% of endogenous retroviruses are found in nearly intact forms, suggesting that these genome regions were acquired relatively recently [13].

In contrast to the abundant retroviral EVEs, examples of extensive non-retroviral EVEs spanning several kilobases also exist, such as the human herpesvirus 6 integration, which exhibits highly homologous repeat sequences to those of human telomere. This suggests that certain genomic sequences in mammalian genome structure have to be discussed in the context of EVEs [14].

In this study, complete viral genome data files registered in NCBI/RefSeq and GenBank (specifically, the virushostdb data file) were utilized to examine their homology to recently completed human T2T full genome (hT2T) [15]. The human genome reference, such as GRCh38 continually developed by the Genome Reference Consortium (GRC), still contain approximately over 180 Mbp of sequence gaps, including portions of the centromeric, pericentromeric, and subtelomeric regions. The T2T genome successfully complements these previously missing regions, achieving a complete, gap-free human genome reference. We report newly identified regions exhibiting long viral sequence homology, especially in centromeric and pericentromeric regions, which were not detected in previous human genome builds prior to hT2T.

Notably, long sequences corresponding to *Entomopoxvirus* were observed, which had not been reported in the previous studies [1, 2], which primarily focus on the distribution, evolution, and diversity of especially non-retroviral EVEs derived from a wide range of well-characterized families of viruses across vertebrate genomes. The *Poxviridae* family is broadly classified into two subfamilies: *Chordopoxvirinae* (which infects vertebrates) and *Entomopoxvirinae* (which infects insects). *Entomopoxviruses* have large, linear, double-stranded DNA genomes and the ability to infect a wide range of insects. Like other *Poxviruses, Entomopoxviruses* replicate exclusively in the host cytoplasm. However, they are unique in their ability to produce spheroid or spindle-shaped occlusion bodies. In contrast, the *Chordopoxviruses* do not typically form these distinct occlusion structures. Furthermore, the *Entomopoxvirus* genome often contains mobile genetic elements, including transposon-like sequences (Polinton-like), which contribute to genome plasticity and evolution [16].

## Results and Discussion

Using the latest hT2T human genome build, we performed homology analysis with the viralhostdb’s complete viral data file (release 224) via the fasta36 software (e-value threshold set at 1e-25). Among sequences with over 57% identity and lengths exceeding 5 kb, the total of unique viral sequences amounted to 2.71 Mb, representing 0.087% of the hT2T genome. Notably, sequences homologous to viruses in the *Entomopoxvirinae* subfamily were identified in centromeres on chromosome (Chr) 3, 13, 21, 22 and X (Fig. 1) (The locations on the *Entomopoxviruses* with high homology to each human chromosome are shown in Supplemental Data1. Furthermore, the actual FASTA36 raw data for the highly homologous region on human Chr 3 in Supplemental Data 1 is presented in Supplemental Data 2.). Of all viruses, *Entomopoxvirus* exhibited the greatest representation, with sequences totaling 2.42 Mb, accounting for 89.0% of all viral homologous regions. These homologous regions were primarily localized near centromeres on the short arms of Chr 13, 21, and 22, mixed with sequences homologous to other viruses in low gene density regions (as shown on heat maps in Fig. 1), with the exception of Chr 3. Homologous regions on the terminal long arm of Chr 20 and near the centromere on the short arm of the X chromosome corresponded to 5 kb and 8 kb regions homologous to Grapevine fleck virus and Clostridium phage, respectively, spanning intronic regions of the RTEL1 and CCNB3 genes. Other homologous regions on Chr 2, 6, 10, 22, and X were primarily composed of stringent simple repeat regions. Detailed locations of these homologous regions within the human hT2T genome are presented in Supplemental Data 3 (Chr 2, 6, 10, 20, and X) and 4 (Chr 3, 13, 21, 22 corresponding to Fig.2 and Supplemental Data 5-7).

**Fig. 1.**
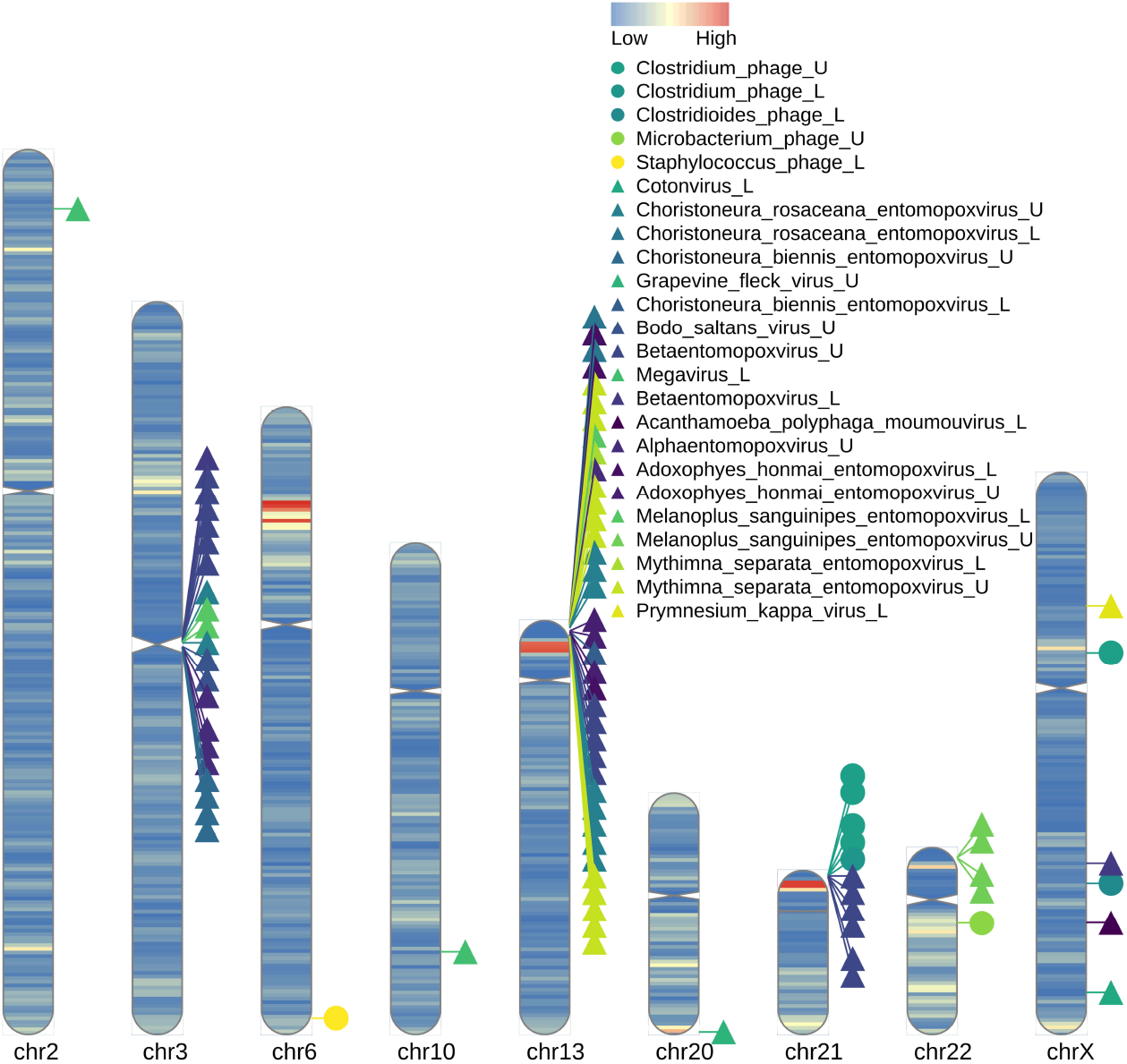
Locations of virus-like sequences and homologous sequences on the human T2T genome. All chromosomes were built using hT2T files (sequence data and .gff annotation file, downloaded from NCBI). Each chromosome is shaded to reflect gene density. Clusters of homologous sequences are observed within the centromere of chromosome 3, and on chromosomes 13, 21, and 22. Most scattered sequences across the genome are tandem simple repeat arrays; however, on chromosome 20, a region with moderate gene density near the end of the long arm contains a sequence homologous to Grapevine fleck virus, forming part of the intronic region of the RTEL1 gene. Additionally, on the X chromosome, a sequence in the proximal short arm near the centromere shows high homology to a Clostridium phage and is located within an intron of the CCNB gene. The suffixes “_L” and “_U” at the end of figure labels indicate sequences with homology to multiple viruses (L) or to a single virus (U). For sequences labeled “_L,” the longest homologous sequence name is provided, and a list of viruses is available in Supplemental data 3 and 4. Chromosomes that did not contain sequences with high homology to viral genomes are not depicted.

To determine whether the DNA-level homology between *Entomopoxviruses* and the human genome was noise originating from low-complexity AT-rich regions, fasta36 was executed using a reference file where the low-complexity regions of the human genome were soft-masked. This revealed that for some regions (*) in Supplemental Data 1, the E2() value significantly exceeded the E() value; therefore, considering low E() value, these were categorized as “Weak signal.” In the other regions, the E2() values were dramatically lower (Supplemental Data 8). It should be noted that when homology searches were performed with both the viral query and the human genome reference soft-masked, almost all signals indicating a link between the two were lost, even when using fasta36. Furthermore, using tfastx36, homology between the full peptide sequences of all viral strains shown in Supplemental Data 1 (queries) and the DNA sequences of the human genomic regions with high DNA-level homology to *Entomopoxviruses* (reference) was searched. This confirmed numerous sequences with E-values of less than 1e-5 in many of the corresponding regions (Supplemental Data 9). Some hits exhibited very low E-values, falling below 1e-10. All data maintained a higher value for functional similarity (Positives) compared to amino acid identity, suggesting functional conservation. Similarly, calculations were performed using the corresponding BLAST family programs (tblastn and tblastx), but these yielded almost no output data. This suggests that for sequences presumed to have interacted in the distant past, as in this case, the determination of common seed sequences becomes very difficult due to the accumulation of mutations. Taken together, the homology between *Entomopoxviruses* and parts of the human genome detected by fasta36 was determined not to be due to noise or chance, but rather to represent a reliable signal of the relationship between the two.

The EVE sequence in the centromere of Chr 3 spanned 748 kb in total, representing 16.25% of the centromeric region (Fig. 2). EVEs on Chr 13, Chr 21, and Chr 22 spanned totally 1.43 Mb (Supplemental Data 5), 291 kb (Supplemental Data 6), and 141 kb (Supplemental Data 7), respectively (See explanation in Supplemental Data 10). These sequences predominantly consist of the satellite DNA hsat1A. No centromeric satellite sequences were found outside of these regions, and hsat1A regions accounted for 96.6% of the total 2.71 Mb viral-homologous regions found in this study. These regions are unique to the hT2T build, with no liftover to GRCh37 or GRCh38 possible and sparse annotation. Analysis of Simple Repeats (table: hub_170992_simpleRepeat) downloaded from the UCSC Genome Browser indicated lower repeat density in other regions compared to Chr 3’s centromere.

**Fig. 2.**
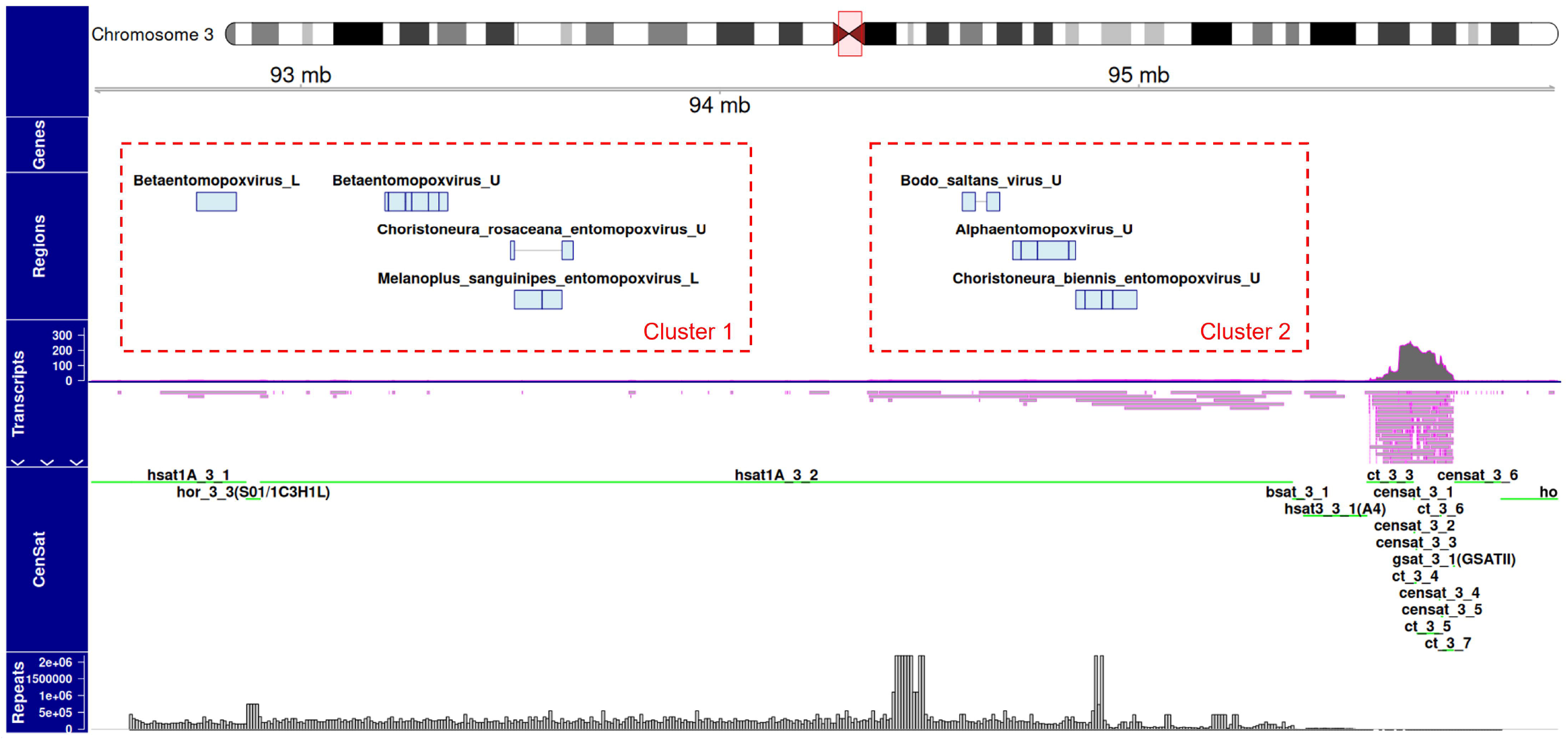
Centromeric region with high similarity to viruses within chromosome 3. This region is unique to hT2T, cannot currently be lifted over from GRCh37 or 38, and shows no distribution of genes. On the left, the “Regions” panel displays bars indicating viral regions with high similarity. The mapping results of long-read RNA-seq data from European human samples used in this study, aligned with minimap2, are shown on the left under “Transcripts.” Although transcription from viral-related regions is observed (bottom of “Transcripts”), transcription levels are low (top of “Transcripts”). Long non-coding RNAs of over 10 kb have been identified in the virus-like sequences. In contrast, numerous transcripts have been observed primarily from the hsat3 region on the right side of the figure. “Repeats” shows the distribution of simple repeats; this region contains numerous intermittent short repeats, displayed here with a bin size of 5,000 and window size of 500. Specifics of all viral regions in Fig.2 are shown in Supplemental Data 4. The integration loci inferred as results of independent events are indicated by red dashed boxes.

While *Entomopoxvirus*-like sequences were identified in the complete human genome, their relatively low identity (a minimum 57%) and the functional significance to core eukaryotic cell division suggest the possibility that ancestral *Entomopoxviruses* may have infected more primitive eukaryotes, leading to eventual endogenization. To investigate, we retrieved homologous sequences from the 249-39,967 bp region of NC_023426 *Alphaentomopoxvirus* anomala cuprea virus, which shares 58% identity with a continuous 37,759 bp region in the hT2T chromosome 3 centromere (Fig. 2a; *Alphaentomopoxvirus*_U box on human Hsat1A_3_2), across 341 eukaryotic species from yeast to mammals (Ensembl release-113), aligning them using FASTA36 (e-value 1e-25). For comparison, homologous sequences to the 249-39,967 bp region in hT2T chromosome 3 were also retrieved across 341 species and aligned with MAFFT software. However, alignments revealed no significant conservation across species for either viral or human homologous regions (data not shown). Given the relatively low identity (minimum 57%) of the sequences identified, the current set of publicly available *Entomopoxvirus* genomic sequences may not include the most closely related homologs of the anciently fixed elements found in the human genome. This considerable evolutionary distance and the potential lack of immediate relatives in the current database significantly complicate the accurate inference of the phylogenetic history and ancestral state of these long virus-like sequences. The low conservation observed in the cross-species alignment is therefore a predictable outcome of both the sequences’ antiquity and the limited scope of the available comparative data,highlighting the inherent uncertainty in performing deep ancestral reconstruction for such highly diverged elements.

As shown in Supplemental Data 1, seven *Entomopoxvirus* species were fragmented into regions less than 40 kb each, with nearly all of these segments scattered in the human genome [17]. Given that *Entomopoxviruses* are linear double-stranded DNA viruses that replicate within the host’s cytoplasm, the mechanism by which these sequences were integrated into the host genome requires discussion. Traditional molecular mechanisms for integration are inadequate to explain this. A plausible hypothesis is that *Entomopoxvirus* sequences might have been integrated during co-infection with retroviruses or by utilizing enzymes by endogenous retroviruses, facilitating chromosomal integration during cell division. Given the low identity (57–70%) and the unlikely scenario of *Entomopoxvirus* infection to primates, such a scenario may have occurred in ancient times, though definitive evidence remains elusive. Furthermore, the possibility remains that the present *Entomopoxvirus* genomic region was acquired partly through horizontal gene transfer from an ancient host. In the present study, we detected transcripts originating from *Entomopoxvirus*-like sequences within the human genome (Fig.2, Supplemental Data 5-7). The presence of these transcripts is evocative of viruses in the family *Poxviridae* that acquired genes from the host genome via Long Interspersed Nuclear Element-1 (LINE-1) [18]. Moreover, recent studies have demonstrated active horizontal DNA transfer among *Entomopoxviruses*, their associated Polinton-Like Viruses (PLVs), and host cells. Specifically, PLVs possess the ability to integrate into the cellular genome as proviruses, and EPV and PLV undergo mosaic genomic evolution through the exchange of gene modules and functional domains [16]. Considering these complex mechanisms of gene acquisition and exchange collectively, the possibility remains that *Entomopoxviruses* utilized these intricate molecular mechanisms, specifically, LINE-1, the PLV-mediated shuttling function, and its mosaic restructuring capability, to construct longer sequences from partial sequences acquired from an ancient host genome and subsequently incorporated them into its own genome, or vice versa. Importantly, few sequence homologous to *Chordopoxvirus* was detected in the human genome in this study. This finding is consistent with the lack of known association between PLVs and the *Chordopoxvirinae* subfamily.

To evaluate whether the integrations of the long virus-like sequences concentrated on Chr 3, Chr 13, Chr 21, and Chr 22 (Fig. 1) occurred as independent events within the genome, the number of integration loci was determined. First, a threshold for defining a locus was set. Fragment groups constituting a single locus were assumed to have resulted from the fragmentation of a single copy of a viral genome through local deletions or rearrangements mediated by host genomic mechanisms. This assumption is based on the known mechanism where full-length integrated retroviral EVEs undergo deletion via homologous recombination between the long terminal repeats (LTRs), often leaving behind a solo LTR [19]. Thus, assuming a full-length integration of a viral genome, viral fragments observed across regions longer than the full-length were considered to be integrations that occurred as independent events. Specifically, since the full length of the *Entomopoxvirus* is approximately 300 kb at its longest, viral fragment clusters present in regions farther apart than this distance were considered to have resulted from independent integration events. Based on this criterion, two independent loci were identified on Chr 3, five on Chr 13, and one each on Chr 21 and Chr 22. This suggests that the integrations separated into Clusters 1-9 occurred as multiple independent events.

The discovery of moderately homologous yet extensive (hundreds of kb) continuous viral-homologous regions in the human genome, other than human herpesvirus-related telomeric associations [14], is unprecedented. These sequences, undetected in GRCh37 or GRCh38, were identified only with hT2T. Remarkably, 96.6% of the viral-homologous 2.71 Mb was comprised of hsat1A. Located in α-satellite non-coding regions of the genome forming centromeres and pericentromeres [20], α-satellite sequences, while they are often assumed to be identical across centromeres, actually show substantial complexity and chromosomal specificity in variations and polymorphisms. Higher-order repeat units (HOR) and Hsat constituting α-satellites feature consecutive repeat sequences but exhibit diversity in monomer types, mutations, and arrays. For example, HOR identity across non-homologous human chromosomes ranges from 50–70% [21] and varies significantly within the same species.

Over the past 20 years, extensive efforts have been devoted to human genome sequencing and assembly (Latest human genome GRCh38 by Genome Reference Consortium), yet an estimated 5-10% of the human genome has remained unexplored “black box” regions [22]. Most of these regions are found within or around the centromeres of chromosomes, containing highly repetitive DNA sequences that complicate assembly from short DNA sequencing reads. However, the recent completion of a telomere-to-telomere human genome assembly (hT2T-CHM13) has allowed a comprehensive characterization of the centromeric and pericentromeric repeats, which make up 6.2% (189.9 Mb) of the genome [15]. Despite this achievement, it is still not possible to conduct molecular phylogenetic analyses on α-satellite regions in non-human species. High-order repeats (HORs), which form the core of the centromere and are essential for kinetochore function and cell division in eukaryotes [20], are believed to contain sequences widely conserved across eukaryotes.

Nonetheless, they cannot be analyzed with current methods. Similarly, Hsat, another class of repeats, plays a crucial role in chromosomal stability, heterochromatin formation, and epigenetic control essential for life [21], but cross-species analyses are not feasible at this time. In fact, a search for homologous sequences across 341 species in this study yielded no regions with significant homology across species, highlighting the need for complete genome assemblies and solid annotations like human T2T level in additional species.

Since the release of the hT2T genome build, functional studies on hsat1A in humans have begun to progress, with transcripts from hsat1A regions identified, and expression differences between normal and tumor cells reported [23]. In this study, we used long-read RNA-seq data from humans and mapped it to the hT2T genome using Minimap2 to examine transcriptional products in regions identified with entomopox-like sequences on Chr 3, 13, 21, and 22. Transcripts were analyzed for read species (bottom panel in “Transcripts” on Fig. 2) and read count (top panel) in Chr 3, 13, and 21, with limited read counts observed in the individual analyzed (European ancestory). A relatively large number of reads were detected primarily in the Hsat3 region on Chr 3, on the other hand, the limited transcripts identified on viral-like sequences (“Regions” in Fig. 2) include long non-coding RNAs spanning several hundred kilobases. Even at low expression levels, a few copies of these transcripts might be sufficient to regulate centromere function on these chromosomes, suggesting they could be functionally relevant molecules.

*Entomopoxvirus*-like sequences identified herein were found in the classical satellite DNA regions, HSat1A and HSat3. Classical satellite sequences have been shown not to be transcriptionally inert, and as research advances concerning the transcripts of HSat2 and HSat3 (satellite non-coding RNAs), their essential status in multiple cellular contexts, including heterochromatin formation and regulation, stress response, and cancer, has been recognized [23]. HSat1A and Hsat3 are also transcribed, similar to other classical satellites, suggesting a transcriptional abnormality in cancer. This transcriptional dysregulation in the pericentromeric region is a general characteristic of tumor cells, and pericentromere transcription itself is associated with cancer. It has also been suggested that the HSat1A/HSat3 region may be a vulnerable pericentromeric locus originally associated with genomic instability. The incorporation of a foreign sequence into this region may be causing, or conversely exploiting, transcriptional abnormalities in these non-coding regions and chromosomal aberrations that are linked to the progression of cancer. Therefore, the entomopoxvirus-like sequences identified in this study raise a point for necessary future functional validation: whether they are involved in the organization of the host’s genome architecture and the regulation of gene expression under pathological conditions, potentially via the HSat1A/HSat3 satellite non-coding RNAs.

## Materials & Methods

All viral genome data were downloaded from https://www.genome.jp/virushostdb/ (release 224), and the full-length human genome hT2T-CHM13v2.0 was obtained from NCBI. A homology search was performed using the entire viral dataset as the query and hT2T as the reference, using fasta36 with parameters -b 1 -d 1 -E 1e-25 -m BB [24]. Preliminary experiments established an identity threshold of 57% identity and 5 kb continuity to ensure complete capture of *Entomopoxvirus* data, which guided human sequence collection (fasta36_data_collection.py). All python codes described here are stored in GitHub repository (https://github.com/ehondo2020/Takoyaki). Using the output files, human chromosomal locations of viral sequences were extracted (human_genomic_position_extract.py), merged to remove duplicates, and the largest range was determined (chromosomal_range_determination.py). The resulting human genomic position data was converted into BED files, and sequentially processed with make_single_range_1.py, _2.py, and _3.py to finalize the list of human genome positions corresponding to each virus for visualization with RIdeogram [25] and Gviz [26].

CenSat and simple_repeats data from the UCSC Genome Browser were downloaded for Gviz visualization. Simple_repeats counts were collected using the options, a bin size of 5,000 and window size of 500. RNA-seq data were obtained from the Whole Genome Sequencing Project and mapped to hT2T-CHM13v2.0 using Minimap2 [27].

Chromosome 3 centromeric regions in the human hT2T genome (37,759 bp) were compared for homology with other eukaryotic species. Genome data for 341 species were downloaded from Ensembl’s FTP site (release-113), and a batch homology search was performed using fasta36 with parameters -b 1 -d 1 -E 1e-25 -m BB.

To validate the homology between *Entomopoxviruses* and the human genome (specifically against low-complexity regions), a dust mask was applied to the human T2T genome (Chr 3, 13, 21, 22, and X) using the pydustmasker library [28]. The queries were the nucleotide sequences of all boxes in Supplemental Data 1. The fasta36 options were set as follows: fasta36 -S -z 21. These options ensured that low-complexity regions were ignored in the soft-masked human T2T genome reference, and statistical validation was secured through base sequence shuffling. tfastx36 was performed using the full peptide sequences of all viral strains listed in Supplemental Data 1 as queries (downloaded from NCBI Virus). The reference utilized the DNA sequences (approx.2MB) of the human T2T genomic regions (Chr 3, 13, 21, 22, and X) that showed high DNA-level homology with *Entomopoxviruses*. (For the purpose of ensuring statistical stringency and proper evaluation of background noise, and to facilitate the distinction of the target regions on T2T Chr 3, 13, 21, 22, and X as well as to expand the search space, the full lengths of human T2T Chr 1 and 4 were added to the reference. After tfastx36 calculation, homologous regions (data) against Chr1 and Chr4 were eliminated (Supplmental Data 9). The threshold was set to -E 1e-5, and other parameters were set to default. Both blastn and tblastn were executed separately with low-complexity filtering enabled and disabled, and the threshold for tblastn was set to -E 1e-5.

## Acknowledgements

Computations were performed on the NIG supercomputer at ROIS National Institute of Genetics, Japan.

## Data Availability

All data, including those pertaining to computational processes, are provided within the manuscript, with the link to code hosted in a GitHub repository, as well as in the supplementary information files. Additional data related to computational processes can be made available upon reasonable request to the corresponding author.

