## Supplementary figures and images for "Entomopoxvirus-like long DNA sequences in human centromeric and peri-centromeric regions"

### Supplemental Data 1

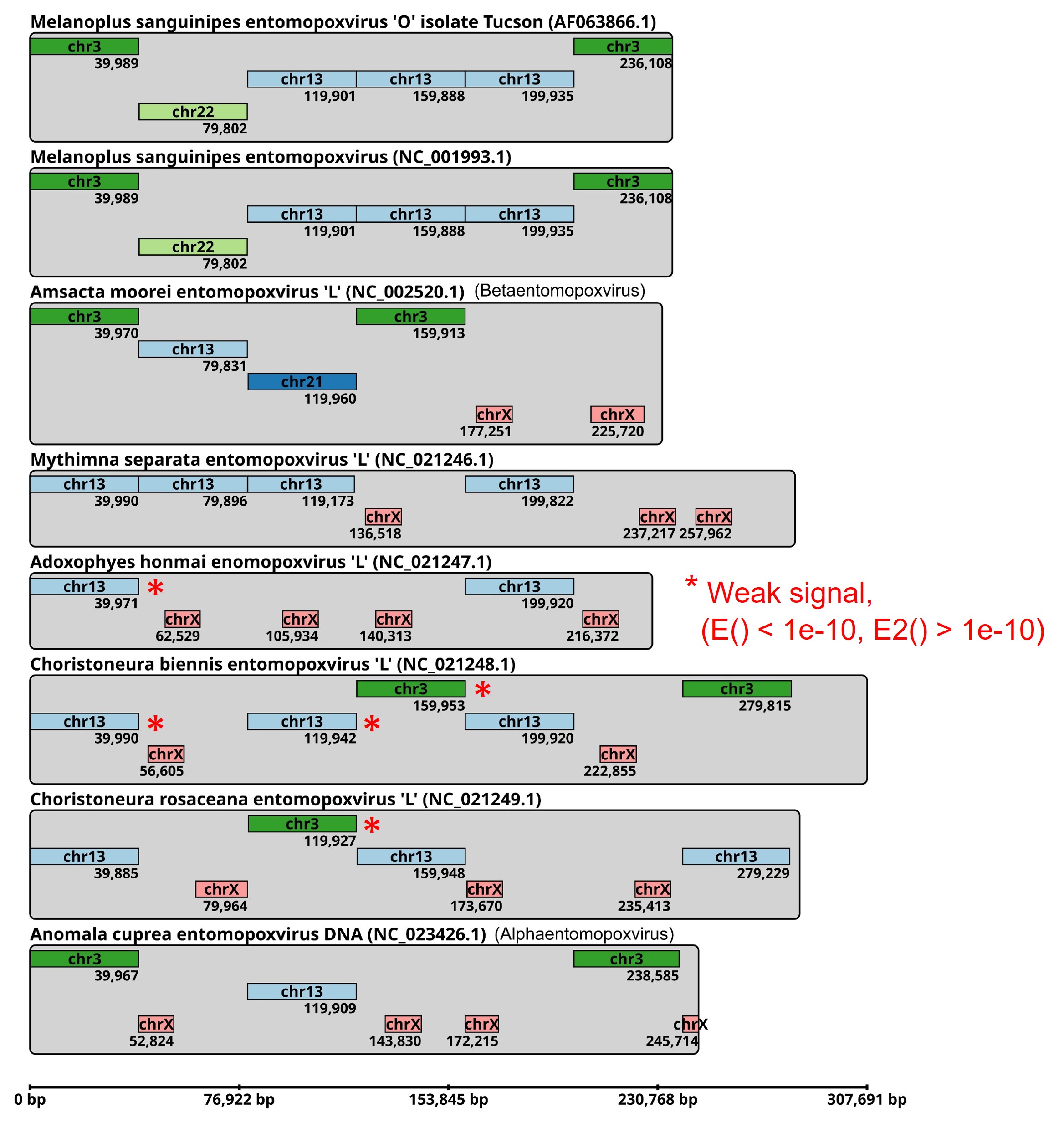

### Supplemental Data 5

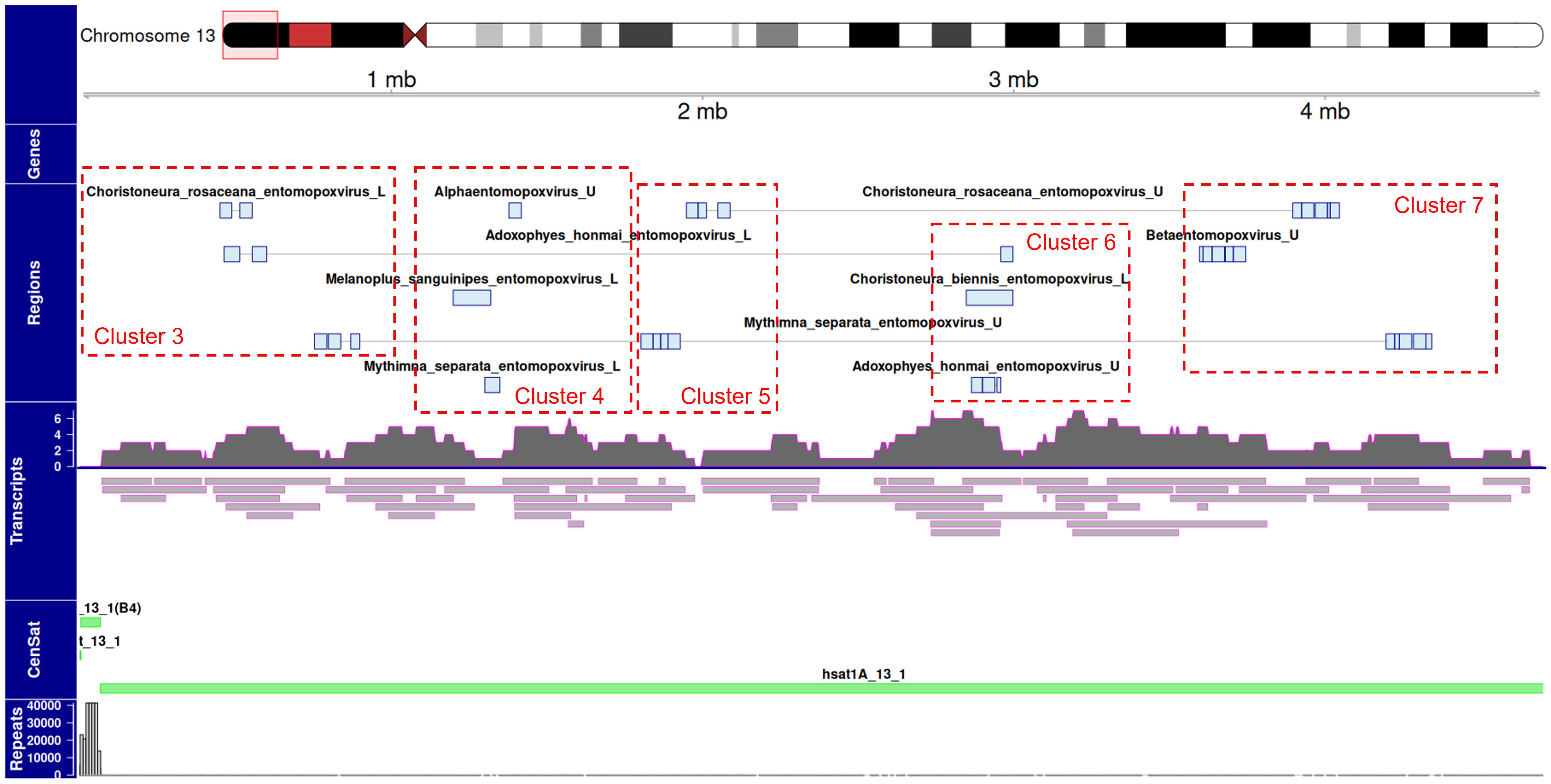

### Supplemental Data 6

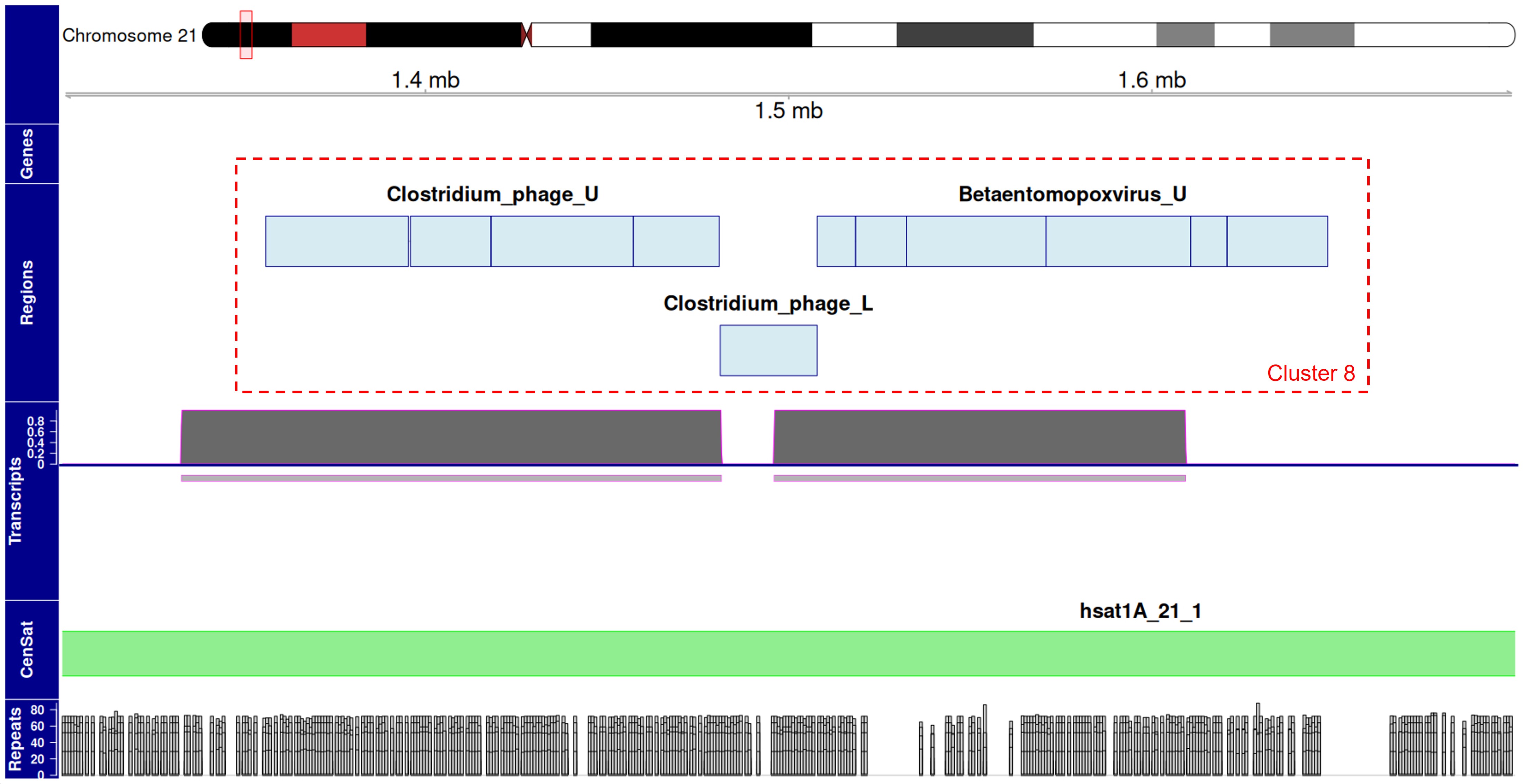

### Supplemental Data 7

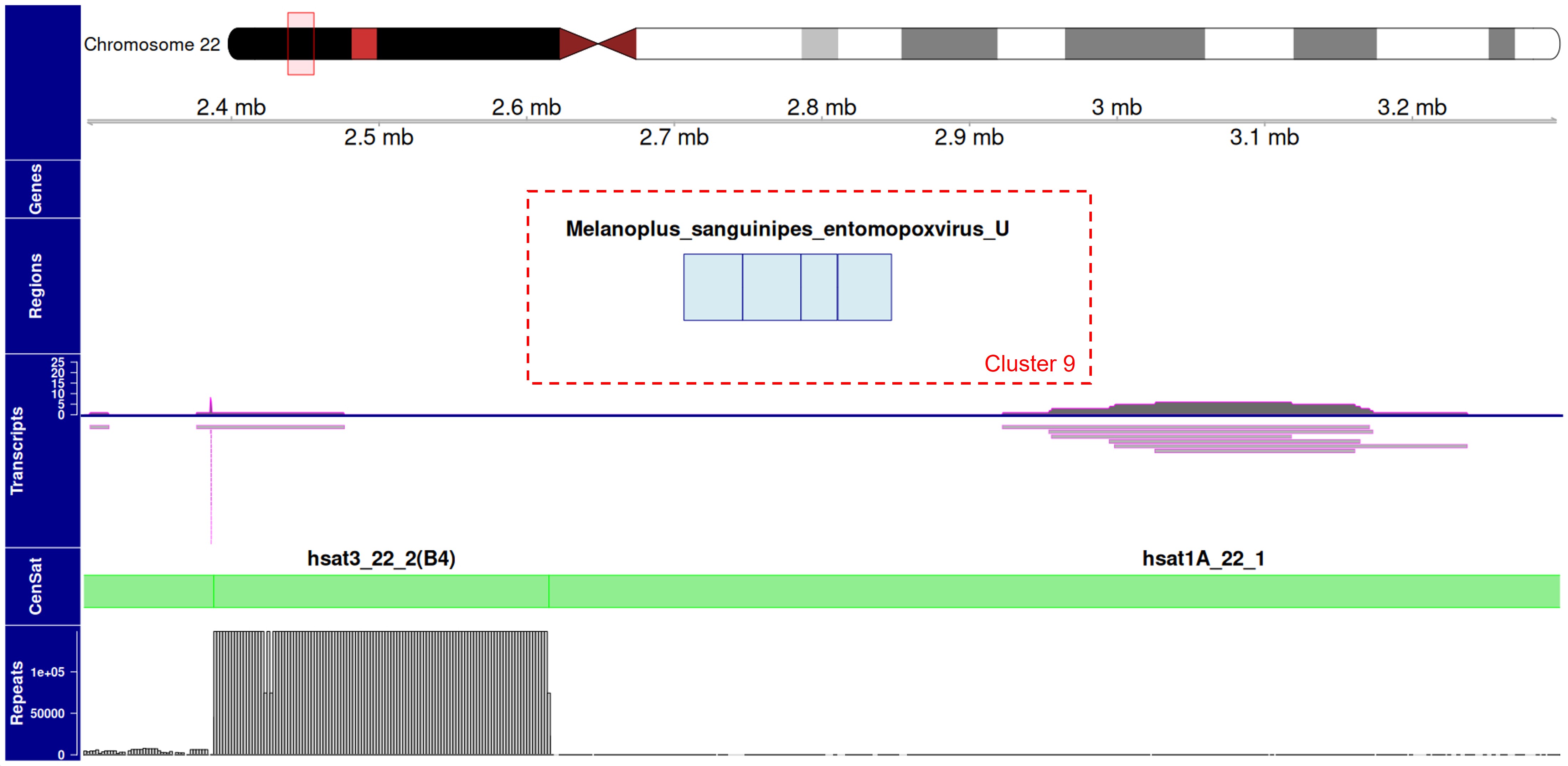
