## Supplemental Data 10 for "Entomopoxvirus-like long DNA sequences in human centromeric and peri-centromeric regions"

################################################################################
# Supplemental Data 10
################################################################################


Figure Legends for Supplemental Data 5-7


Supplemental Data 5  Virus-like sequences distributed in the short arm of chromosome 13. Non-coding RNAs are transcribed across this entire region, with transcripts observed from all parts of the virus-like sequences. Simple repeats are scarcely detected. Shown with a bin size of 5,000 and window size of 500. Specifics of all viral regions in Supplemental Data 5 are shown in Supplemental Data 4. The integration loci inferred as results of independent events are indicated by red dashed boxes.

Supplemental Data 6  Virus-like sequences distributed in the short arm of chromosome 21. This region includes one unique sequence each for entomopoxvirus and Clostridium phage (unlike "_L," these sequences do not overlap with other viral sequences). The small Clostridium phage box (Clostridium_phage_L) contains sequences overlapping with other viruses. Intriguingly, the mapping of transcripts reveals non-coding RNA sequences over 100 kb in length, which closely match the positions of the two virus-unique sequences. Simple repeats are also sparse in this region. Displayed with a bin size of 5,000 and window size of 500. Specifics of all viral regions in Supplemental Data 6 are shown in Supplemental Data 4. The integration loci inferred as results of independent events are indicated by red dashed boxes.

Supplemental Data 7  Virus-like sequences distributed in the short arm of chromosome 22. Only a unique sequence for entomopoxvirus is identified in this region; however, no transcripts are detected, and no particularly distinctive features are observed. Specifics of all viral regions in Supplemental Data 7 are shown in Supplemental Data 4. The integration loci inferred as results of independent events are indicated by red dashed boxes.
